# RAPID ALKALINIZATION FACTOR 22 is a key modulator of the proliferation and hyper-elongation responses of root hairs to microbial volatiles in Arabidopsis

**DOI:** 10.1101/2023.07.20.549818

**Authors:** Rafael Jorge León Morcillo, Jesús Leal-López, Lidia López-Serrano, Edurne Baroja-Fernández, Samuel Gámez-Arcas, Verónica G. Doblas, Alberto Férez-Gómez, Javier Pozueta-Romero

## Abstract

RAPID ALKALINIZATION FACTOR (RALF) peptides are important players in regulating cell expansion. In Arabidopsis, volatile compounds (VCs) emitted by the fungal phytopathogen *Penicillium aurantiogriseum* promote root hair (RH) proliferation and hyper-elongation through ethylene and enhanced photosynthesis signalling actions. A striking alteration in the proteome of fungal VC-treated roots involves up-regulation of RALF22. To test the possible involvement of RALF22 in the fungal VC-promoted RH changes, we characterized RH density and number responses to fungal VCs in *ralf22* and *fer-4* plants impaired in RALF22 and its receptor FERONIA, respectively. Unlike WT plants, *ralf22* and *fer-4* RHs responded weakly to fungal VCs, strongly indicating that the RALF22-FERONIA module is a key determinant of the RH response to fungal VCs. To investigate the regulatory mechanisms behind this response, we analysed the *RALF22* transcript levels in roots of *etr1-3* and *eir1* ethylene signalling mutants and those of ethylene-responsive, RH-related *RSL4, RHD2, PRX1* and *PRX44* transcripts in *ralf22* and *fer-4* roots. Moreover, we characterized the RH and *RALF22* transcript level responses to fungal VCs of the *cfbp1* mutant defective in photosynthetic responsiveness to VCs. Unlike in WT roots, fungal VCs weakly enhanced *RALF22* expression in *etr1-3, eir1* and *cfbp1* roots, and *RSL4, RHD2, PRX1* and *PRX44* expression in *ralf22* and *fer-4* roots. In addition, fungal VCs weakly promoted RH changes in *cfbp1* roots. Collectively, our findings showed that the ethylene and enhanced photosynthesis signalling-mediated RH response to fungal VCs involves RALF22-FERONIA.

## INTRODUCTION

Secreted, small (ca. 5 kDa) RAPID ALKALINIZATION FACTOR (RALF) peptide hormones have emerged as important players in multiple procesess, including root and root hair (RH) growth and development, response and tolerance to biotic and abiotic stresses, regulation of plant-microbe interactions and modulation of immune responses (Atkinson et al. 2013, Bergonci et al. 2014, Du et al. 2016, Murphy et al. 2016, Stegmann et al. 2017, Zhao et al. 2018, Blackburn et al. 2020, Zhu et al. 2020, Abarca et al. 2021, Tang et al. 2022). In Arabidopsis, RALF peptides comprise a family of 37 members and practically all of them have the same function of regulating cell expansion (do Canto et al. 2014). The majority of these small peptides are ligands of plasma membrane-localized protein complexes involving FERONIA (Zhao et al. 2018, Abarca et al. 2021), a receptor-like kinase that regulates growth, development and responses to environmental changes through mechanisms involving calcium signalling, alkalinization of the extracellular medium, interaction with hormones and the production of reactive oxygen species (ROS) (Deslauriers and Larsen 2010, Yu et al. 2012, Haruta et al. 2014, Chen et al. 2016, Du et al. 2016, Feng et al. 2018, Zhu et al. 2020, Martínez-Pacheco et al. 2023). There are numerous studies analyzing aspects of this complex signalling network (Blackburn et al. 2020, Abarca et al. 2021). However, the functional role of the majority of RALF peptides is largely unknown. Most previous studies have been based on the use of very young (4-7 days-after sowing) seedlings cultured *in vitro* in sucrose containing medium (Srivastava et al. 2009, Atkinson et al. 2013, Haruta et al. 2014, Du et al. 2016, Zhao et al. 2018, Zhu et al. 2020, Abarca et al. 2021, Li et al. 2022, Liu et al. 2023). Amongst all investigated Arabidopsis RALF peptides, only RALF1 has been shown to regulate RH development in very young seedlings (Du et al. 2016, Zhu et al. 2020). Although lateral roots (LRs) are major determinants of the ability of the root system to acquire water and nutrients from the soil, all the studies on the involvement of the RALF1-FERONIA signalling module on RH growth and development have been centred on the primary root of very young seedlings (Duan et al. 2010, Du et al. 2016, Zhu et al. 2020).

In nature, plants communicate with microorganisms by exchanging chemical signals through the phytosphere (Huang et al. 2019, Singh et al. 2023). Such interactions are important for plant productivity, fitness and capacity to adapt to environmental changes (De-la-Peña and Loyola-Vargas 2014, Stringlis et al. 2018, He et al. 2022). In the precolonization phase, before direct contact with plants occurs, microorganisms emit a large number of volatile compounds (VCs) that promote plant growth, nutrient acquisition and photosynthesis (Zhang et al. 2008, Zhang et al. 2009, Sánchez-López et al. 2016, García-Gómez et al. 2020, Gámez-Arcas et al. 2022). These compounds may cause massive LR formation and RH growth, thus improving the root‟s capacity to explore for water and minerals and host microorganisms (Gutiérrez-Luna et al. 2010, Ditengou et al. 2015, Garnica-Vergara et al. 2016, García-Gómez et al. 2019, García-Gómez et al. 2020). We have recently shown that VCs from *Penicillium aurantiogriseum* (a fungal phytopathogen that can be found in the rhizosphere [Bodini et al. 2011, Gharaei-Fathabad et al. 2014; Kłapeć et al. 2018]) enhance plant growth and photosynthesis, and promote changes in root system architecture (RSA), including proliferation and shortening of LRs, and proliferation and “hyper-elongation” of RHs on LRs (García-Gómez et al. 2019, 2020). These morphological changes were found to involve ethylene and enhanced photosynthesis signalling and were associated with up-regulated expression of ROS (e.g. O_2_^-^ and H_2_O_2_) producers (e.g. RHD2, PRX1 and PRX44) at both transcript and protein levels, augmented levels of ethylene biosynthetic enzymes (e.g. SAM1, SAM2, ACO2 and CAS-C1) and enhanced ethylene emissions by roots (García-Gómez et al. 2020).

A striking change in the proteome of roots of fungal VC-treated plants involved strong up-regulation of RALF22 (At3g05490) (García-Gómez et al. 2020). RALF22 accumulates mainly in roots (https://pax-db.org/protein/619670) and participates with FERONIA in the transduction of cell-wall signals to regulate plant growth and salt stress tolerance (Zhao et al. 2018). To investigate the possible involvement of the RALF22-FERONIA complex in the RSA changes promoted by fungal VCs, we characterized the LR and RH responses to fungal VCs of *ralf22* and *fer-4* plants impaired in RALF22 and FERONIA expression, respectively. Furthermore, to investigate the possible involvement of the RALF22-FERONIA complex in the ethylene-mediated RH response to fungal VCs, we characterized the response to fungal VCs of *RALF22* transcripts in roots of ethylene-signalling mutants and that of ethylene-responsive *RSL4, RHD2, PRX1* and *PRX44* transcripts involved in RH formation and elongation in *ralf22* and *fer-4* roots. Moreover, to obtain knowledge on the mechanisms of enhanced photosynthesis signalling that modulate fungal VC-promoted enhancement of RALF22 expression and RH proliferation and elongation, we characterized the responses to fungal VCs of RHs and *RALF22* transcript levels in roots of the *cfbp1* mutant defective in photosynthetic responsiveness to VCs (Ameztoy et al. 2021). Our findings establish the central role of RALF22-FERONIA in the ethylene and enhanced photosynthesis signalling-mediated RH response to fungal VCs. A possible involvement of RALF22-FERONIA in microbe-induced mechanisms for hijacking the plant ethylene signalling pathway is discussed.

## RESULTS

### RALF22 and FERONIA are required for the fungal VC-promoted proliferation and hyper-elongation responses of RHs on LRs

In the absence of small fungal VCs, *ralf1, ralf22* and *fer-4* plants exhibited similar to WT growth (**Figure 1A,B**) and LR number and length phenotypes (**Figure 1C**). RHs on LRs of *fer-4* plants were shorter and less abundant than on WT LRs, whereas those of *ralf22* plants exhibited a size and abundance similar to WT (**Figure 1C**). Like in WT plants, fungal VCs promoted rosette and root growth in WT, *ralf1* and *ralf22* plants, although their effect in *fer-4* plants was weaker than in WT plants (**Figure 1A,B**). Fungal VCs also promoted LR proliferation and shortening similar to the WT in *ralf1, ralf22* and *fer-4* plants (**Figure 1C, Supplemental Figure 1**), strongly indicating that RALF1, RALF22 and/or FERONIA do not mediate the LR response to fungal VCs. Notably, unlike WT and *ralf1* plants, *ralf22* and *fer-4* plants were unresponsive in terms of VC-promoted proliferation and elongation of RHs on LRs (**Figure 1C**, **Supplemental Figure 1**).

**Figure 1.**
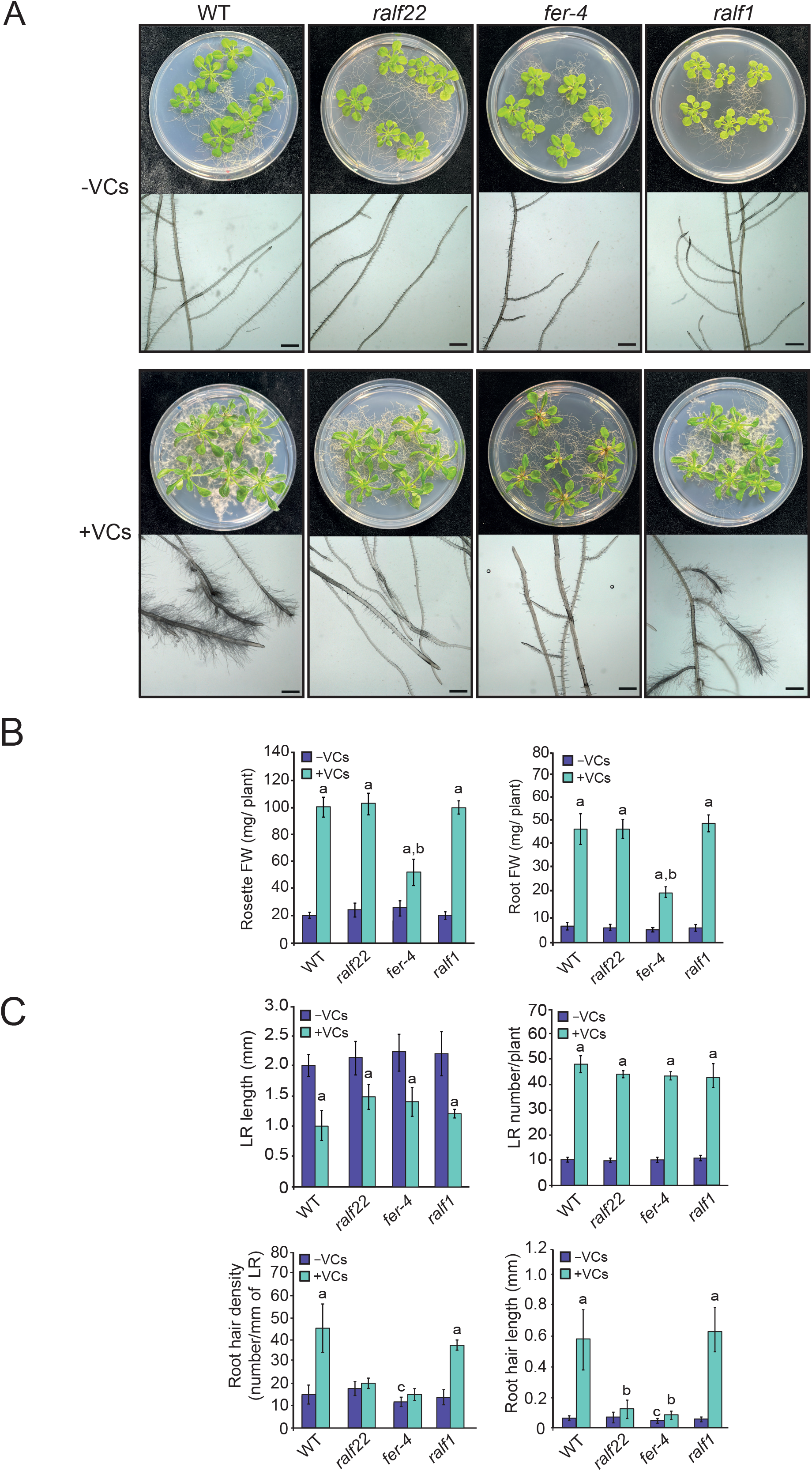
RALF22 and FERONIA are required for the fungal VC-promoted proliferation and hyper-elongation responses of RHs on LRs. (A) External phenotypes, (B) rosette and root FW and (C) root architecture parameters of WT, *ralf22, ralf1* and *fer-4* plants cultured in the absence or continuous presence of fungal VCs for one week. Values in panels (B) and (C) are means ± SE for three biological replicates (each a pool of 12 plants) obtained from four independent experiments. Letters “a” and “b” indicate significant differences, according to Student‟s t-test (P<0.05), between: “a” VC-treated and non-treated plants, “b” VC-treated WT and mutants and “c” WT plants and mutants cultured without fungal VC treatment. RH number and length data were obtained from a pool of 6 LRs per plant. Plants providing data shown in (A) and (B) were grown on horizontal plates, whereas those providing data in (C) were grown on vertical plates. Scale bars in (A), 1 mm.

### RALF22 mediates the ethylene-regulated RH response to fungal VCs

Fungal VC-promoted RH proliferation and hyper-elongation involves ethylene perception and signalling (García-Gómez et al. 2020). These responses are associated with enhanced ethylene emissions by roots and up-regulation of the expression of some ethylene biosynthetic enzymes, including CAS-C1 and ACO2, at both transcript and protein levels (García-Gómez et al. 2020). To investigate the possible involvement of RALF22 in the ethylene-mediated RH response to fungal VCs, we analysed the *RALF22* transcript level responses in the roots of two mutants whose RHs weakly respond to fungal VCs (García-Gómez et al. 2020): the *etr1-3* ethylene receptor and the *eir1* ethylene insensitive mutants. In addition, we analysed by qRT-PCR the levels of transcripts encoding proteins involved in ethylene perception (e.g. *ETR1, ERS1* and *ERS2*), signalling (e.g. *EIN2, EIN3* and *CTR1*) and synthesis (e.g. *CAS-C1* and *ACO2*) in roots of WT and *ralf22* plants cultured with/without fungal VCs. Furthermore, we characterized the RH response of WT and *ralf22* plants to exogenous application of 1-aminocyclopropane 1- carboxylate (ACC), the substrate of the enzyme that catalyses the last step of the ethylene biosynthetic pathway. These analyses revealed that, unlike in WT roots, fungal VCs did not enhance *RALF22* transcript levels in *etr1-3* and *eir1* roots (**Figure 2A, Supplemental Figure 2**). Fungal VCs enhanced the transcript levels of *ERS2* in WT roots, but not in *ralf22* roots (**Figure 2B**), and promoted changes similar to those in the WT in the expression of *ETR1, ERS1, EIN2, EIN3, CTR1, CAS-C1* and *ACO2* in *ralf22* roots (**Supplemental Figure 2**). Moreover, ACC application promoted RH proliferation and elongation in WT plants, but not in *ralf22* plants (**Figure 2C**).

**Figure 2.**
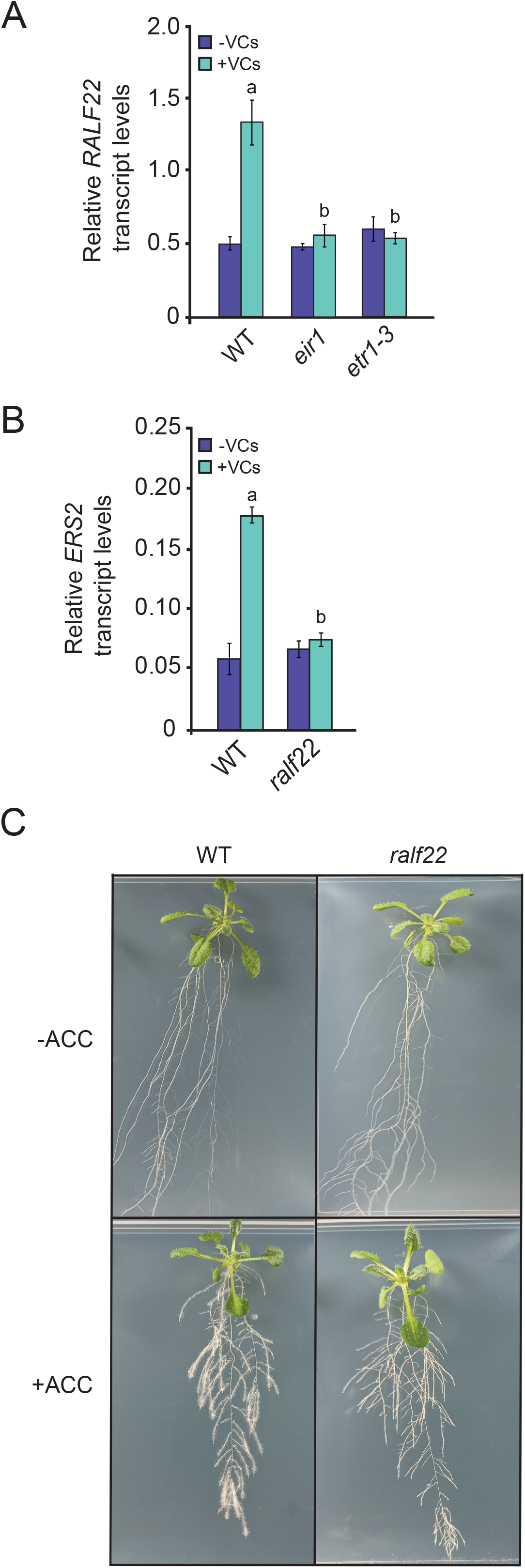
RALF22 mediates the ethylene-regulated RH response to fungal VCs. (A) Relative abundance of *RALF22* transcripts as determined by qRT-PCR in roots of WT, *eir1* and *etr1-3* plants cultured on vertical plates in the absence or continuous presence for one week of fungal VCs. (B) Relative abundance of *ERS2* transcripts in roots of WT and *ralf22* plants cultured on vertical plates in the absence or continuous presence for one week of fungal VCs. (C) External phenotypes of WT and *ralf22* plants grown on vertical plates with or without 1 μm ACC supplementation in the culture medium. Values in (A) and (B) are means ± SE for three biological replicates (each a pool of 4 plants) obtained from three independent experiments. Letters “a” and “b” indicate significant differences, according to Student‟s t-test (P<0.05), between: “a” VC-treated and non-treated plants, and “b” VC-treated WT and mutants.

### The RALF22-FERONIA complex mediates the fungal VC-promoted up-regulation of ROS producers

The RH response to fungal VCs is a ROS (e.g. O_2_^-^ and H_2_O_2_)-dependent process that is associated with enhanced activities of the NADPH oxidase RHD2 and class III apoplastic peroxidases (e.g. PRX1 and PRX44) (García-Gómez et al. 2020). These functions are required for RH growth (Foreman et al. 2003, Martin et al. 2022, Marzol et al. 2022) and are regulated by the RH length-determining RSL4 transcription factor (Yi et al. 2010, Mangano et al. 2017), which in turn is regulated by ethylene (Song et al. 2016, Feng et al. 2017) and the FERONIA-activated early translation initiation factor eIF4E1 (Zhu et al. 2020). We hypothesized that the response of RHs to fungal VCs involves, at least in part, RALF22-FERONIA complex-mediated up-regulation of *RSL4, RHD2, PRX1* and *PRX44* expression. In line with this presumption, qRT-PCR analyses revealed that fungal VCs strongly enhanced the levels of *RSL4, RHD2, PRX1* and *PRX44* transcripts in WT roots, but not in *ralf22* and *fer-4* roots (**Figure 3**).

**Figure 3.**
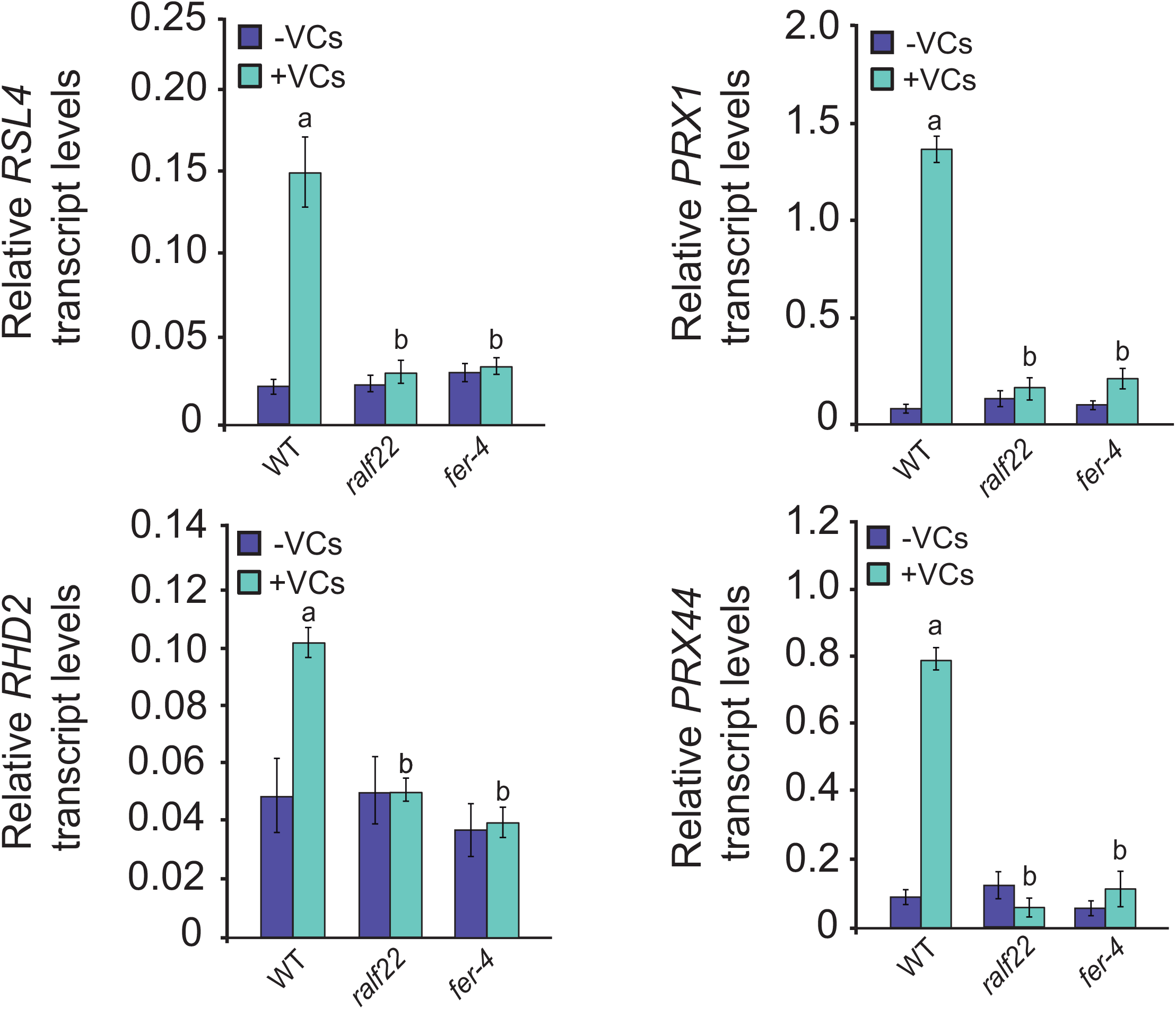
The RALF22-FERONIA complex mediates the fungal VC-promoted up-regulation of ROS producers. Relative abundance of *RSL4, RHD2, PRX1* and *PRX44* transcripts as determined by qRT-PCR in roots of WT, *ralf22* and *fer-4* plants cultured in the absence or presence for one week of fungal VCs. Values are means ± SE for three biological replicates (each a pool of 4 plants) obtained from three independent experiments. Letters “a” and “b” indicate significant differences, according to Student‟s t-test (P<0.05), between: “a” VC-treated and non-treated plants, and “b” VC-treated WT and mutants.

### RALF22-FERONIA-mediated RH responses to fungal VCs are subject to enhanced photosynthesis signalling

Soluble sugars photosynthetically produced in leaves act as hormone-like signalling molecules that are transported to roots to support and adjust growth and development according to the shoot demand (Lemoine et al. 2013, Tong et al. 2022). Fungal VCs cause an increase in photosynthetic activity and soluble sugar content (Sánchez-López et al. 2016, García-Gómez et al. 2019). Because exogenously supplied sugars promote RH proliferation and elongation (Jain et al. 2007, Mishra et al. 2009, Karve et al. 2012), we hypothesized that the RALF22- FERONIA-mediated RH responses to fungal VCs are due, at least in part, to enhanced photosynthesis signalling mediated by sugars. To test this hypothesis, we used qRT-PCR to analyse the levels of *RALF22* transcripts in roots of WT, *fer-4* and *cfbp1* plants defective in photosynthetic responsiveness cultured with or without supplementary sucrose in the absence or presence of fungal VCs. Moreover, we characterized the RH proliferation and elongation responses of these plants to fungal VCs.

Without sucrose supplementation, fungal VC exposure strongly enhanced *RALF22* transcript levels in WT roots, but did so weakly in *cfbp1* roots (**Figure 4**). Sucrose supplementation enhanced *RALF22* expression in roots of WT and *cfbp1* plants, especially when exposed to fungal VCs (**Figure 4**). Notably, fungal VCs did not affect *RALF22* expression in roots of *fer-4* plants cultured in either the absence or presence of exogenously supplied sucrose (**Figure 4**), strongly indicating that the *RALF22* expression response to fungal VCs is subject to control by FERONIA. Sucrose application slightly promoted RH elongation in both WT and *cfbp1* roots (**Figure 5**). Without sucrose supplementation, the RH proliferation and elongation responses to fungal VCs of *cfbp1* plants were weaker than those of WT plants (**Figure 5**). Notably, unlike in *ralf22* and *fer-4* plants, fungal VCs strongly promoted the proliferation and elongation of RHs in *cfbp1* plants cultured in sucrose-containing medium (**Figure 5**, **Supplemental Figure 3**).

**Figure 4.**
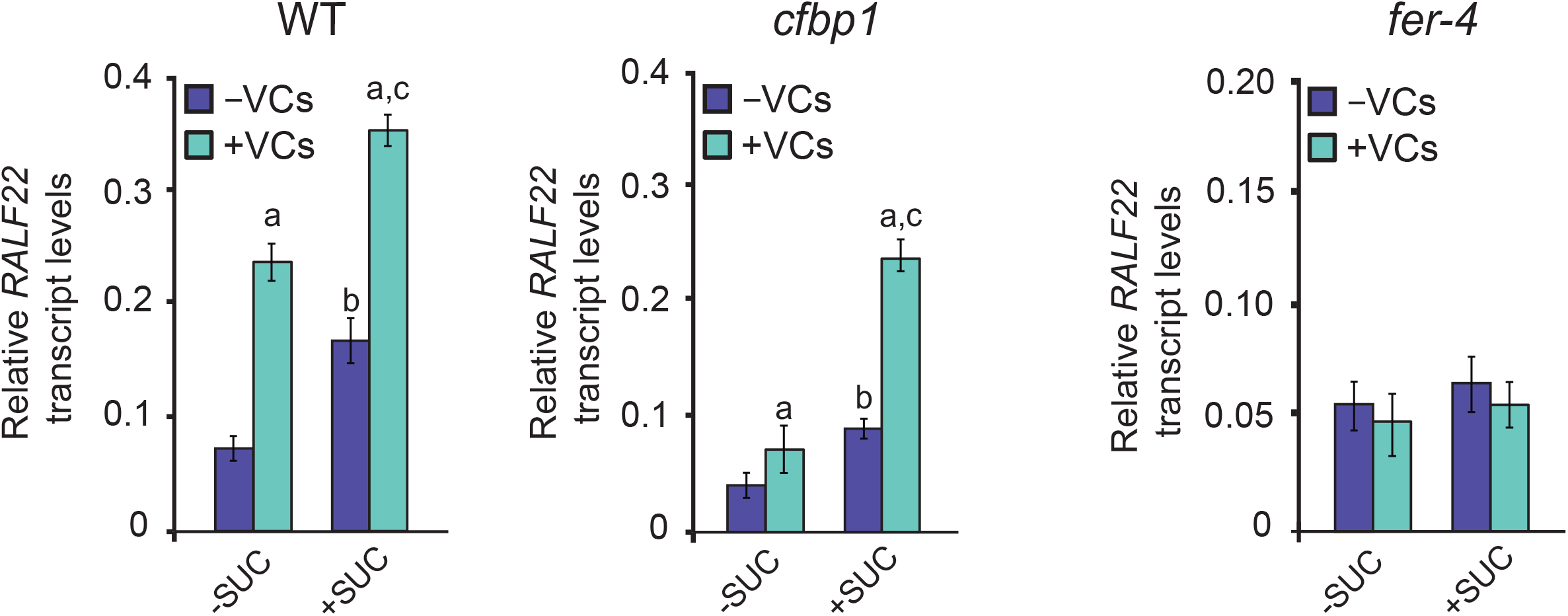
The RALF22 expression response to fungal VCs is subject to sugar-mediated enhanced photosynthesis signalling. Relative abundance of *RALF22* transcripts as determined by qRT-PCR in roots of WT, *cfbp1* and *fer-4* plants cultured in the absence or continuous presence for one week of fungal VCs, with or without 45 mM sucrose supplementation in the culture medium. Values are means ± SE for three biological replicates (each a pool of 4 plants) obtained from three independent experiments. Letters “a”, “b” and “c” indicate significant differences, according to Student‟s t-test (P<0.05), between: “a” VC-treated and non-treated plants, “b” sucrose-treated and non-treated plants and “c” VC-treated plants cultured with or without 45 mM sucrose supplementation.

**Figure 5.**
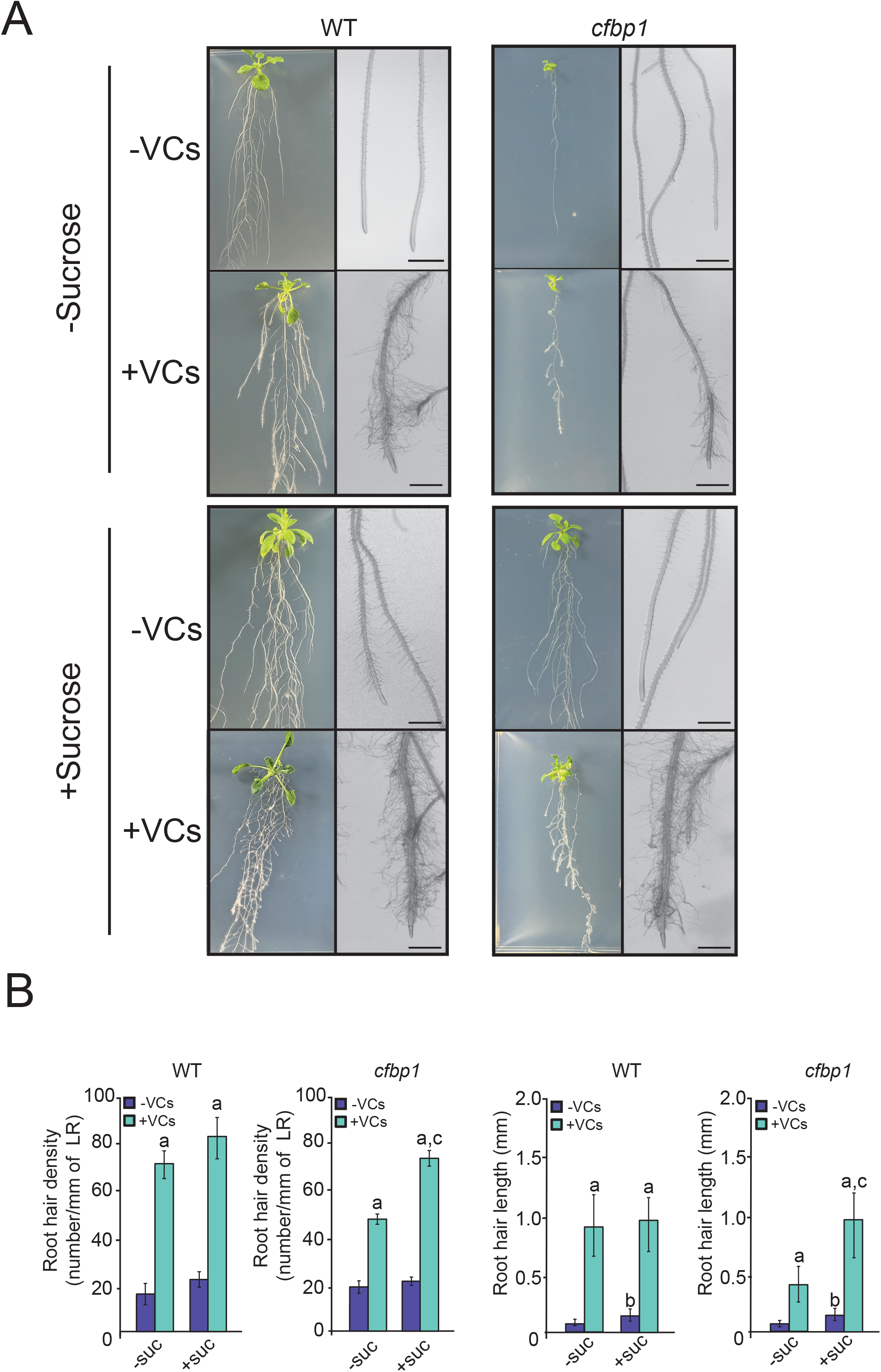
RALF22-FERONIA-mediated RH responses to fungal VCs are subject to enhanced sugar-mediated photosynthesis signalling. (A) External phenotypes and (B) RH parameters (density and length) of WT and *cfbp1* plants cultured on vertical plates in the absence or continuous presence of fungal VCs for one week, with or without 45 mM sucrose supplementation in the culture medium. Values in panel (B) are means ± SE for three biological replicates (each a pool of 12 plants) obtained from four independent experiments. Letters “a”, “b” and “c” indicate significant differences, according to Student‟s t-test (P<0.05), between: “a” VC-treated and non-treated plants, “b” sucrose-treated and non-treated plants and “c” VC-treated plants cultured with or without sucrose supplementation. Scale bars in (A), 1 mm.

### *Penicillium aurantiogriseum* emits ethylene

Some fungal phytopathogens including *Penicillium* spp. produce ethylene which, once perceived by the plant, leads to an ethylene burst response (Marcos et al. 2005, Yang et al. 2017, Ballester and González-Candelas 2020). Several factors strongly indicated that the RALF22-FERONIA-dependent RH response to *P. aurantiogriseum* VCs could be triggered, at least partly, by fungal ethylene emissions, namely: (i) some root morphological changes promoted by fungal VCs, including root shortening and RH elongation, are similar to those of plants exposed to exogenous ethylene (Pitts et al. 1998, Negi et al. 2008, Street et al. 2015) (**Figure 1**); (ii) fungal VCs enhance the expression of ethylene-inducible genes including *RSL4, PRX1, PRX44, RHD2, ERS1, ERS2, CAS-C1* and *ACO2* (**Figure 2B**, **Figure 3, Supplemental Figure 2**) and the accumulation of ROS in RHs (García-Gómez et al. 2020); (iii) RSA and *RALF22* expression in roots of the *etr1-3* mutant impaired in the ethylene ETR1 receptor weakly responded to *P. aurantiogriseum* VCs (García-Gómez et al. 2020) (**Figure 2**); and (iv) the promoter region of *RALF22* contains several cis-elements for key transcription factors of the ethylene signalling pathway including EIN3 and ORA59 (Chang et al. 2013, Yang et al. 2021). To test this hypothesis, we analysed the ethylene content in the headspace of growth chambers containing *P. aurantiogriseum* cultures. These analyses revealed substantially higher levels of ethylene in the headspace of growth chambers containing the fungal cultures than in controls (0.006 ± 0.002 and 0.972 ± 0.120 ppm, respectively).

## DISCUSSION

By greatly increasing the absorptive surface area, RHs play important roles in acquisition of nutrients and water, plant anchorage and microbe interactions (Grierson et al. 2014, Verbon et al. 2023). Their formation and elongation are genetically defined and environmentally regulated by factors including nutrients and hormones and their interactions with sugars. This work sheds light on the mechanisms underlying RH elongation promoted by ethylene-containing fungal volatile emissions and demonstrates that, in LRs of adult plants, the ethylene and enhanced photosynthesis signalling-mediated RH responses to fungal VCs involve RALF22-FERONIA. To our knowledge, this work provides the first evidence for the involvement of RALF peptides in the elongation of RHs in response to biotic triggers. Because RALF22 positively regulates RH elongation but does not exert any negative effect on LR number and length, modulation of RALF22 expression in RHs has a considerable application potential to improve nutrient and water acquisition and thus crop yield. Therefore, investigating plant responses to microbial VCs is an excellent way not only to understand plant-microbe interactions, but also to uncover fundamental mechanisms involved in developmental regulation in plants and to obtain valuable clues for crop improvement.

### RALF22, but not RALF1, is involved in the fungal VC-promoted RH proliferation and elongation responses in LRs

RALF1 cooperatively acts with FERONIA to regulate RH development in primary roots of young seedlings (Du et al. 2016, Zhu et al. 2020). Here we found that, unlike in RHs of LRs of *ralf22* plants, fungal VCs promoted proliferation and hyper-elongation of RHs on LRs of *ralf1* plants similar to those of WT plants. Thus, RALF22, but not RALF1, is an important determinant of the fungal VC-promoted proliferation and hyper-elongation responses of RHs on LRs. These findings were surprising because (i) exogenous application of different RALFs promote RH elongation (Gjetting et al. 2020, also confirmed in our lab), and (ii) it has been predicted that RALF1 and RALF22 have similar expression patterns (Cao and Shi, 2012). However, unlike RALF22, RALF1 expression in roots is not enhanced by fungal VCs (García-Gómez et al. 2020). Further expression analyses at the tissue and cellular levels will be necessary to understand how RALF peptides regulate RH formation and how they orchestrate their actions in response to environmental changes to prevent redundancy.

In the context of seedling and root growth, FERONIA is involved in the perception and/or signalling pathway of the majority of the RALF peptides (Abarca et al. 2021). Here we found that *fer-4* mutant plants exhibited a similar to *ralf22* unresponsiveness of RHs to fungal VCs (**Figure 1**), which strongly indicates that the RALF22-dependent RH response to fungal VCs involves FERONIA. Notably, fungal VCs did not enhance *RALF22* transcript levels in *fer-4* plants (**Figure 4**). It thus appears that, in addition to its role in the perception and/or signalling of RALF22, FERONIA may play a role in fine-tuning of transcriptional feedback regulation of *RALF22* expression in response to fungal VCs.

### The RALF22-FERONIA complex is involved in the ethylene signalling-and ROS-mediated RH response to fungal VCs

Some root morphological changes promoted by VCs emitted by *P. aurantiogriseum* are similar to those triggered by exogenous ethylene application (Pitts et al. 1998, Negi et al. 2008, Street et al. 2015). Here we found that *P. aurantiogriseum* emits ethylene. Therefore, it is likely that, in the co-cultivation system used in this study, the RALF22-FERONIA-dependent RH response to fungal VCs is triggered, at least partly, by fungal ethylene emissions. As ethylene and FERONIA generally decrease root branching (Negi et al. 2008, Dong et al. 2018), the increased number of LRs observed in fungal VC-exposed plants (**Figure 1**) suggests that *P. aurantiogriseum* emits bioactive VCs different from ethylene that are transduced through signalling pathways different from FERONIA and that are important determinants of LR and RH formation. In addition to ethylene, *P. aurantiogriseum* also emits carbon monoxide (CO) and nitric oxide (NO) (García-Gómez et al. 2019). The exogenous application of these gaseous hormones promotes LR formation and RH elongation (Guo et al. 2008, Guo et al. 2009, Méndez-Bravo et al. 2010, Lombardo and Lamattina 2012). Therefore, it is likely that the RSA and RH changes promoted by fungal VCs are, at least partly, due to CO and NO emissions.

Ethylene signalling increases ROS accumulation to drive RH initiation and elongation through mechanisms involving RSL4-mediated up-regulation of class III PRXs and NADPH oxidases (Mangano et al. 2017, Martin et al. 2022). Some of these ROS producers (e.g. PRX1, PRX44 and RHD2) are strongly up-regulated by fungal VCs (García-Gómez et al. 2020). RSL4 expression is transcriptionally regulated by ethylene (Song et al. 2016, Feng et al. 2017) and by mechanisms mediated by the RALF1-FERONIA complex that involve enhanced capacity of *RSL4* transcripts to bind to the early translation initiation factor eIF4E1 (Zhu et al. 2020). In a previous work we demonstrated that the RH response to fungal VCs is ROS-dependent, as RHs of RHD2-lacking *rhd2* plants did not respond to fungal VCs (García-Gómez et al. 2022). Here we provided strong evidence that, in the *in vitro* co-cultivation system used in this and previous studies, the ethylene-and ROS-mediated RH proliferation and elongation responses to fungal VCs involve RALF22-FERONIA-RSL4 action, as schematically illustrated in the model shown in **Figure 6**. First, application of ACC and ethylene-containing blends of fungal VCs promoted RH proliferation and elongation in WT plants, but not in *ralf22* and *fer-4* plants (**Figure 1** and **Figure 2C**). Second, fungal VCs up-regulated the levels of ethylene-inducible ROS-and RH-related *RSL4, RHD2, PRX1* and *PRX44* transcripts in roots of WT plants, but not in *ralf22* and *fer-4* roots (**Figure 3**). Third, unlike in WT roots, fungal VCs did not enhance *RALF22* transcript levels in roots of the *etr1-3* and the *eir1* ethylene insensitive mutants (**Figure 2A**). According to this model, the RALF22-FERONIA-RSL4-mediated RH proliferation and elongation response to fungal VCs is initiated by the binding of fungal ethylene to receptors, with ETR1 playing a dominant regulatory role. Downstream ethylene signalling mechanisms transcriptionally up-regulate the expression of genes involved in ethylene biosynthesis (e.g. *SAM1, SAM2* and *ACO2*) and subsequent production of endogenous ethylene, which once perceived by the receptors, further intensify ethylene signalling. In addition, ethylene signalling transcriptionally up-regulates the expression of *RALF22* and *RSL4*. Once translated and exported to the apoplast, RALF22 binds to FERONIA which phosphorylates the eIF4E1 early translation factor, which in turn regulates *RSL4* translation. This transcription factor triggers the transcription of RH-related genes including those encoding RHD2, PRX1 and PRX44 that produce O_2_^-^ and H_2_O_2_ in the apoplast for RH elongation. The positive regulation exerted by RALF22-FERONIA on fungal VC-promoted up-regulation of SAM1 and SAM2 and ethylene production in roots (García-Gómez et al. 2020) apparently conflicts with Mao et al. (2015), who proposed that FERONIA interacts with SAM1 and SAM2 to suppress ethylene biosynthesis.

**Figure 6.**
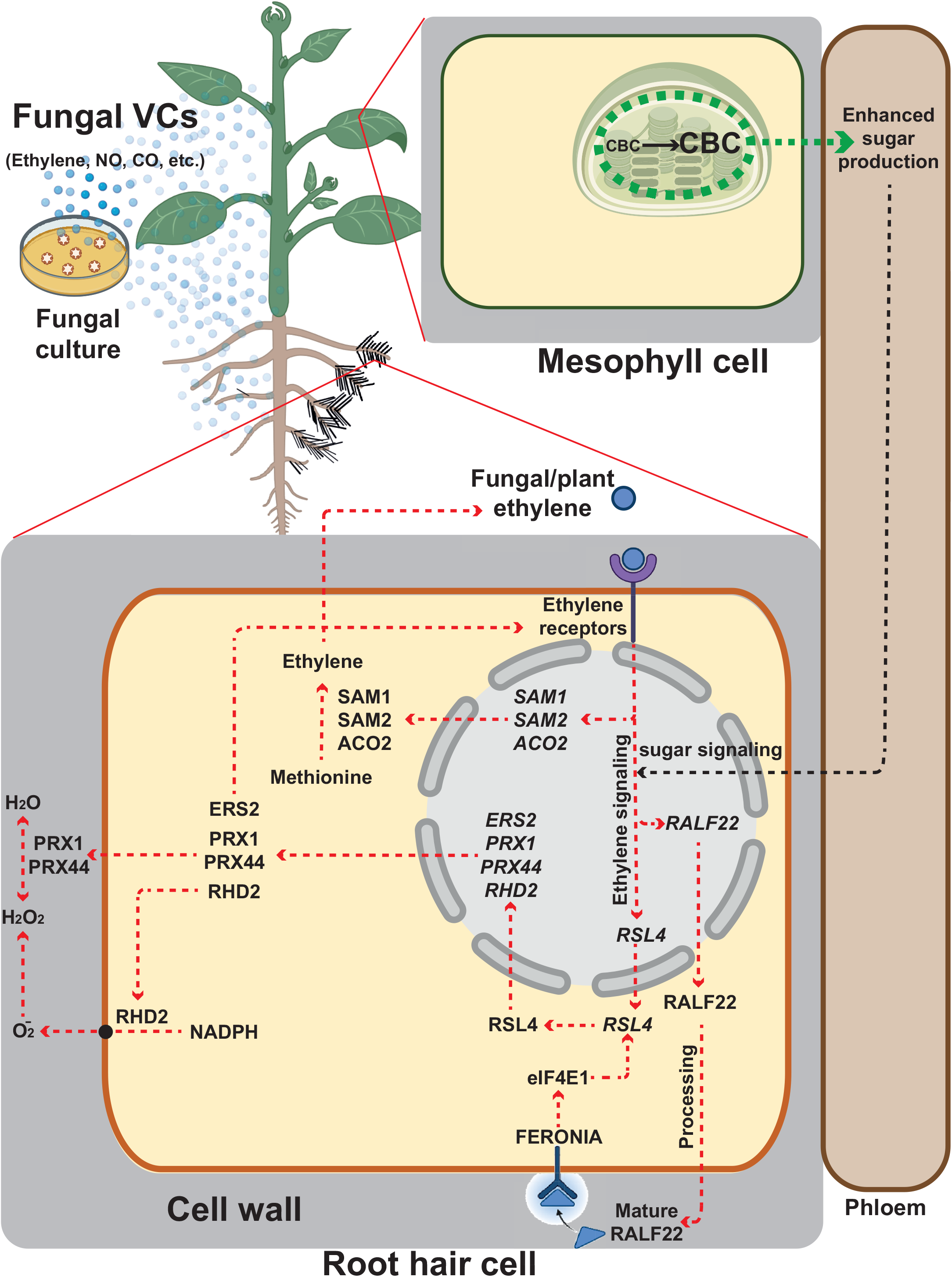
Suggested hypothetical model of regulation of the RH proliferation and hyper-elongation response to fungal VCs by the RALF22-FERONIA complex. According to this model, the response of RHs to fungal VCs involves RALF22-FERONIA-mediated mechanisms wherein long-distance signalling of enhanced photosynthesis and signalling of fungal and endogenously produced ethylene play important roles in the expression of RH-related genes. RH proliferation and elongation response to fungal VCs is initiated by the binding of fungal ethylene to membrane-bound receptors and the incorporation of sugars (mainly sucrose) into the RH. Downstream ethylene signalling mechanisms transcriptionally up-regulate the expression of genes involved in ethylene biosynthesis (e.g. *SAM1, SAM2* and *ACO2*) and the subsequent production of endogenous ethylene, which once perceived by the receptors further intensify ethylene signalling. In addition, ethylene and sucrose signalling transcriptionally up-regulate the expression of *RALF22* and *RSL4*. Once translated and exported to the apoplast, RALF22 binds to FERONIA which phosphorylates the eIF4E1 early translation factor, which in turn regulates the translation of *RSL4*. This transcription factor triggers the transcription of RH genes including those encoding RHD2, PRX1 and PRX44 that produce O ^-^ and H O in the apoplast for RH elongation. Figure was created using Biorender.com.

Ethylene modulates its own synthesis through mechanisms involving ethylene signalling and transcriptional regulation of the expression of ethylene biosynthetic genes, both by positive and negative feedback mechanisms (Pattyn et al. 2021). ERS2 is one member of the ethylene receptor family. Its expression is transcriptionally regulated by ethylene and interacts with ETR1 to activate it (Hua et al. 1998, Gao et al. 2008). The fact that, unlike in WT roots, fungal VCs did not enhance *ERS2* transcript levels in *ralf22* roots (**Figure 2B**) strongly indicated that the ethylene signalling and production in fungal VC-exposed plants is subject to tight regulation of the RALF22-FERONIA pathway (**Figure 6**). It thus appears that, contrary to Deslauriers et al. (2010) who proposed that FERONIA does not form part of the ethylene signalling pathway, RALF22-FERONIA plays an important role in the homeostatic regulation of ethylene synthesis and signalling in fungal VC-exposed roots.

### Sugar-mediated enhanced photosynthesis signalling is an important determinant of the RH response to fungal VCs mediated by RALF22

Sugars act as signalling intermediates in multiple pathways, including those involved in root responses to environmental changes (Freixes et al. 2002). Fungal VCs increase photosynthetic activity and soluble sugar content (Sánchez-López et al. 2016, García-Gómez et al. 2019). Here we provided strong evidence that the RALF22-mediated RH proliferation and hyper-elongation responses to fungal VCs involve, at least partly, sugar-mediated signalling of enhanced photosynthesis. First, fungal VC-promoted enhancement of *RALF22* expression in roots of *cfbp1* plants cultured without sucrose supplementation was much weaker than in roots of WT plants (**Figure 4**). Second, sucrose supplementation enhanced *RALF22* transcript levels in WT and *cfbp1* roots, especially in VC-exposed plants (**Figure 4**). Third, the RH proliferation and elongation responses to fungal VCs of *cfbp1* plants cultured without sucrose were weaker than those of WT plants (**Figure 5**). The weak responses of *cfbp1* RHs to fungal VCs could be reverted to being much more similar to the WT by exogenous sucrose supplementation (**Figure 5**).

In roots of plants not exposed to fungal VCs, sucrose supplementation promoted a strong enhancement of *RALF22* transcript levels (**Figure 4**), but a relatively weak RH proliferation and elongation response (**Figure 5**). This indicated that high RALF22 expression promoted by sugar feeding is not the sole important player in the fungal VC-promoted RH proliferation and hyper-elongation responses. Ethylene is a positive regulator of RH development (Tanimoto et al. 1995, Pitts et al. 1998) and its synthesis is enhanced by sucrose feeding via molecular mechanisms as yet to be identified (Seneweera et al. 2003, Jeong et al. 2010). Under conditions of elevated photosynthetic CO_2_ fixation, ethylene and sugar production increases (Dhawan et al. 1981, Pan et al. 2019, Smet et al. 2020, Ahammed and Li 2022) and RHs elongate (Niu et al. 2011). Plants cultured with both sucrose and ethylene-containing fungal VCs developed more and substantially longer RHs than those of plants cultured with sucrose only (**Figure 5**). Therefore, although sugars and ethylene often act in an antagonistic manner to each other (Yanagisawa et al. 2003, Price et al. 2004, Karve et al. 2012, Tong et al. 2022), we hypothesize that the RALF22-FERNONIA-mediated response of RHs to fungal VCs is due, at least partly, to the synergistic actions of signalling of enhanced photosynthetic production of sugars and of fungal and plant-produced ethylene (**Figure 6**).

### Additional remarks on the possible involvement of RALF22-FERONIA in microbe-induced mechanisms for hijacking the plant ethylene signalling pathway

Ethylene may play an active role in plant resistance to invading fungi or, alternatively, facilitate fungal invasion (van Loon et al. 2006). Some fungal phytopathogens secrete proteins to hijack the ethylene signalling pathway as a decoy strategy to redirect the host metabolism in favor of the fungus (Darino et al. 2021). Others secrete functional homologues of plant RALF peptides to increase infectious potential, suppress host immunity and alter root architecture (Masachis et al. 2016, Thynne et al. 2017). Here we found that *P. aurantiogriseum* can emit ethylene, which combined with photosynthetically produced sugars, up-regulates RALF22 expression to modulate ethylene signalling. We speculate that some fungi emit ethylene as a decoy, RALF22- FERONIA-dependent strategy to hijack the ethylene signaling pathway of plants and thus promote root developmental changes in favor of the fungi.

## MATERIALS AND METHODS

### Plant and microbial cultures, growth conditions and sampling

The experiments were carried out using *Arabidopsis thaliana* L. (Heynh) WT plants (ecotype Col-0), the *ralf22* knockout mutant (Zhao et al. 2018), the *ralf1* mutant (SALK_036331), the *fer-4* mutant (Duan et al. 2010), the *cfbp1* knockout mutant impaired in the redox-sensitive fructose-1,6-bisphosphatase isoform 1 of the Calvin-Benson cycle (Ameztoy et al. 2021), the *etr1-3* ethylene receptor mutant (N3070) (Hua and Meyerowitz, 1998) and the *eir1* ethylene insensitive and auxin efflux carrier mutant (N8058) (Luschnig et al. 1998). Unless otherwise indicated the plants were cultured in Petri dishes (92 x 16 mm, Ref. 82.1472.001, Sarstedt) containing sucrose-free half-strength solid Murashige and Skoog (MS) (Phytotechlab M519) medium in growth chambers providing „long day‟ 16 h light (90 µmol photons sec^-1^ m^-2^), 22 °C /8 h dark, 18 °C cycles. *P. aurantiogriseum* was cultured in small Petri dishes (35 x 10 mm, Sarstedt, Ref. 82.1135.500) containing solid MS medium supplemented with 90 mM sucrose. Effects of microbial VOC-depleted VCs on plants were investigated using the “box-in-box” co-cultivation system, as described in García-Gómez et al. (2019). Briefly, plant cultures 14 days after sowing and fungal cultures in unlidded Petri dishes were placed in sterile plastic boxes (200 x 150 x 50 mm IT200N Instrument Trays; AW Gregory, UK) sealed with polyvinyl chloride plastic wrap (Gámez-Arcas et al. 2022). As negative controls, Petri dishes containing plants were cultured in sealed boxes together with Petri dishes each containing sterile microbial culture media. After the incubation time indicated for each experiment, roots were harvested, immediately freeze-clamped and ground to a fine powder in liquid nitrogen with a pestle and mortar.

### Root morphological analysis

Ten days after sowing plants cultured on vertical square Petri dishes (10 x 10 x 2 cm, Sarstedt, Ref. 82.9923.422) were placed in a vertical position in sealed plastic boxes containing fungal cultures. After 6 days of co-cultivation, the numbers and lengths of the plants‟ roots and RHs were measured using an ZEISS SteREO Discovery.V12 stereomicroscope (Zeiss, Germany). RH lengths were measured on ca. 2 cm long LRs. Photomicrographs were captured with a Axiocam 503 color camera (Zeiss, Germany). RHs were measured in a region of 5 mm from the LR tips.

### Headspace analysis of ethylene content

For measurement of the ethylene contents in sealed growth boxes containing *P. aurantiogriseum* cultures, the sealed growth boxes were carefully drilled with a syringe. Ethylene contained in the air was determined by gas chromatography as described by García et al. (2010).

### Real-time quantitative PCR

Total RNA was extracted from frozen Arabidopsis roots of *in vitro* cultured plants using the Trizol method according to the manufacturer‟s recommendations (Invitrogen), following treatment with RNAase-free DNAase (Takara). RNA (1.5 μg) was reverse-transcribed using polyT primers and an Expand Reverse Transcriptase kit (Roche) according to the manufacturer‟s instructions. RT-PCR amplification was performed using the primers listed in **Supplemental Table 1**, and their specificity was checked by separating the obtained products on 1.8% agarose gels.

### Statistical analysis

Unless otherwise indicated, data presented here are means (± SE) obtained from 3-4 independent experiments, with 3-5 replicates for each experiment. The significance of differences between plants not exposed to VCs, and plants exposed to *P. aurantiogriseum* VCs, and between WT plants and mutants was statistically evaluated with Student‟s t-tests using the SPSS package. Differences were considered significant if P<0.05.

## ACKNOWLEDGEMENTS

This work was supported by the Ministerio de Ciencia e Innovación (MCIN) and Agencia Estatal de Investigación (AEI) / 10.13039/501100011033/ (grant PID2019-104685GB-100).

## AUTHORS CONTRIBUTIONS

R J L M and J P-R designed the experiments and analyzed the data; R J L M, J L-L, L L-S, E B-F, S G-A, V G D and A F-G performed most of the experiments and discussed the data; R J L M, J L-L and J P-R wrote the article with contributions from all the authors; J P-R and R J L M conceived the project and research plans.

## SUPPLEMENTAL DATA

**Supplemental Figure 1.**
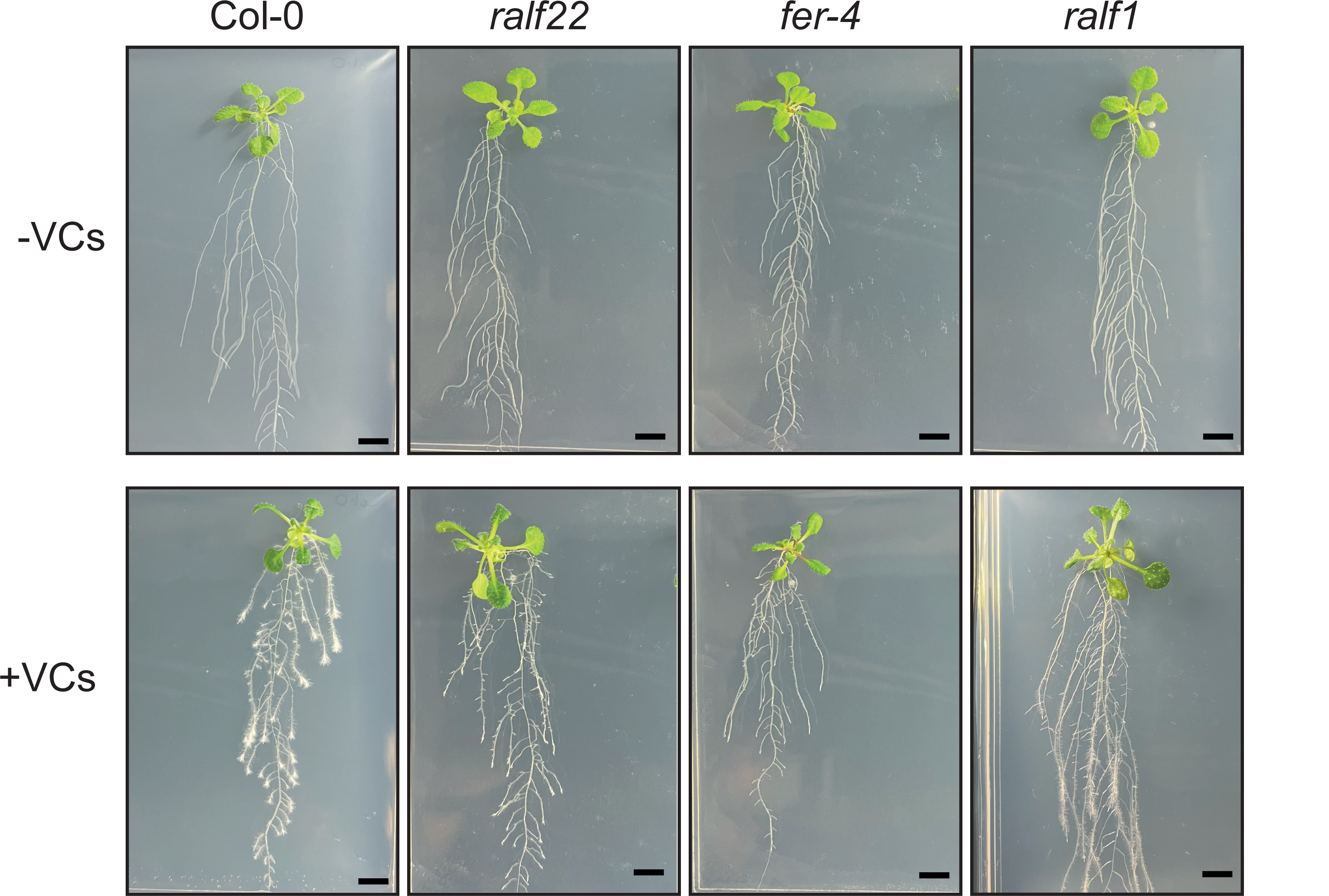
External phenotypes of WT, *ralf22, fer-4* and *ralf1* plants grown on vertical plates in the absence or continuous presence of fungal VCs for one week.

**Supplemental Figure 2.**
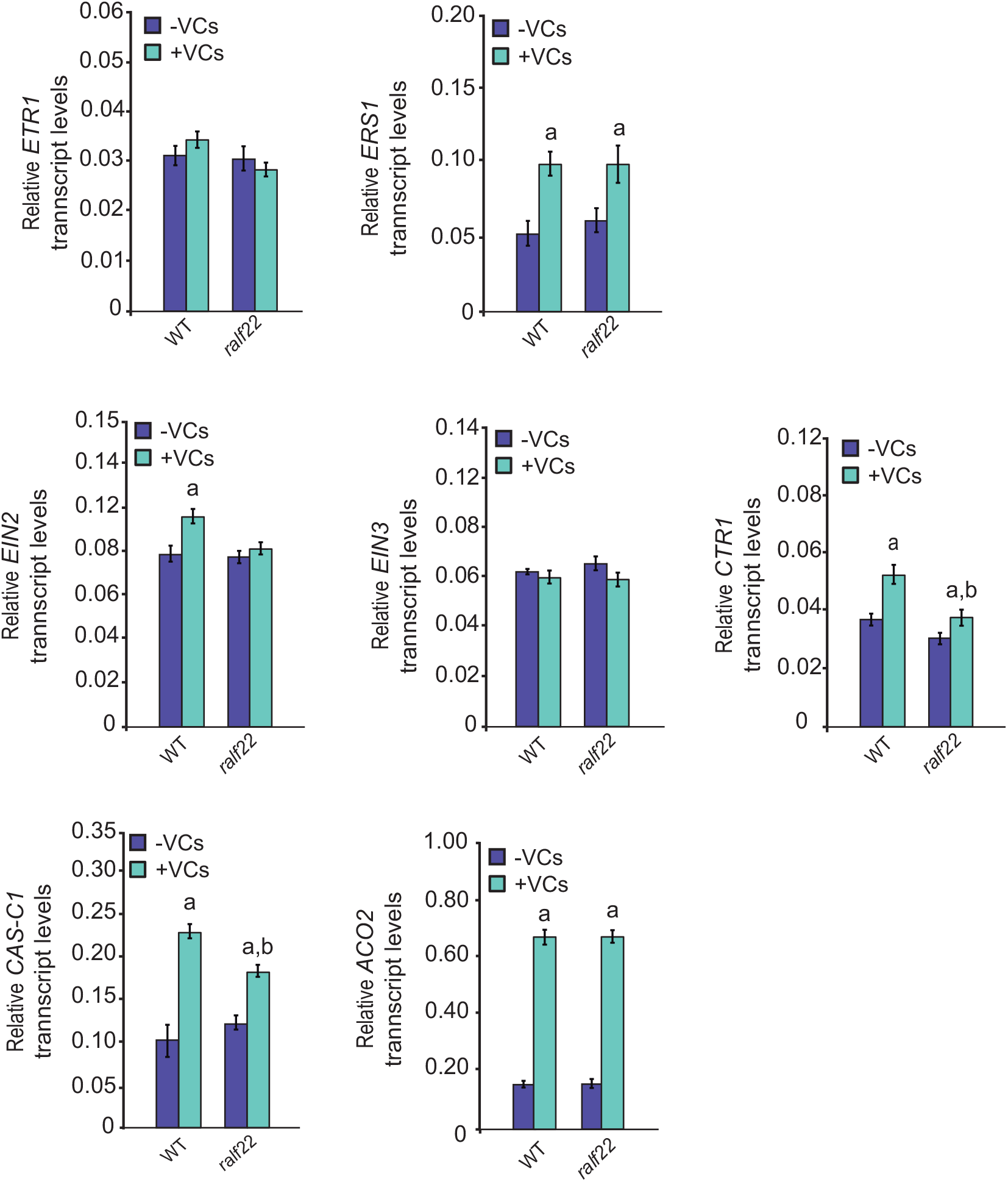
Relative abundance of transcripts involved in ethylene perception, signaling and synthesis in roots of WT and *ralf22* plants cultured in the absence or presence for one week of fungal VCs.

**Supplemental Figure 3.**
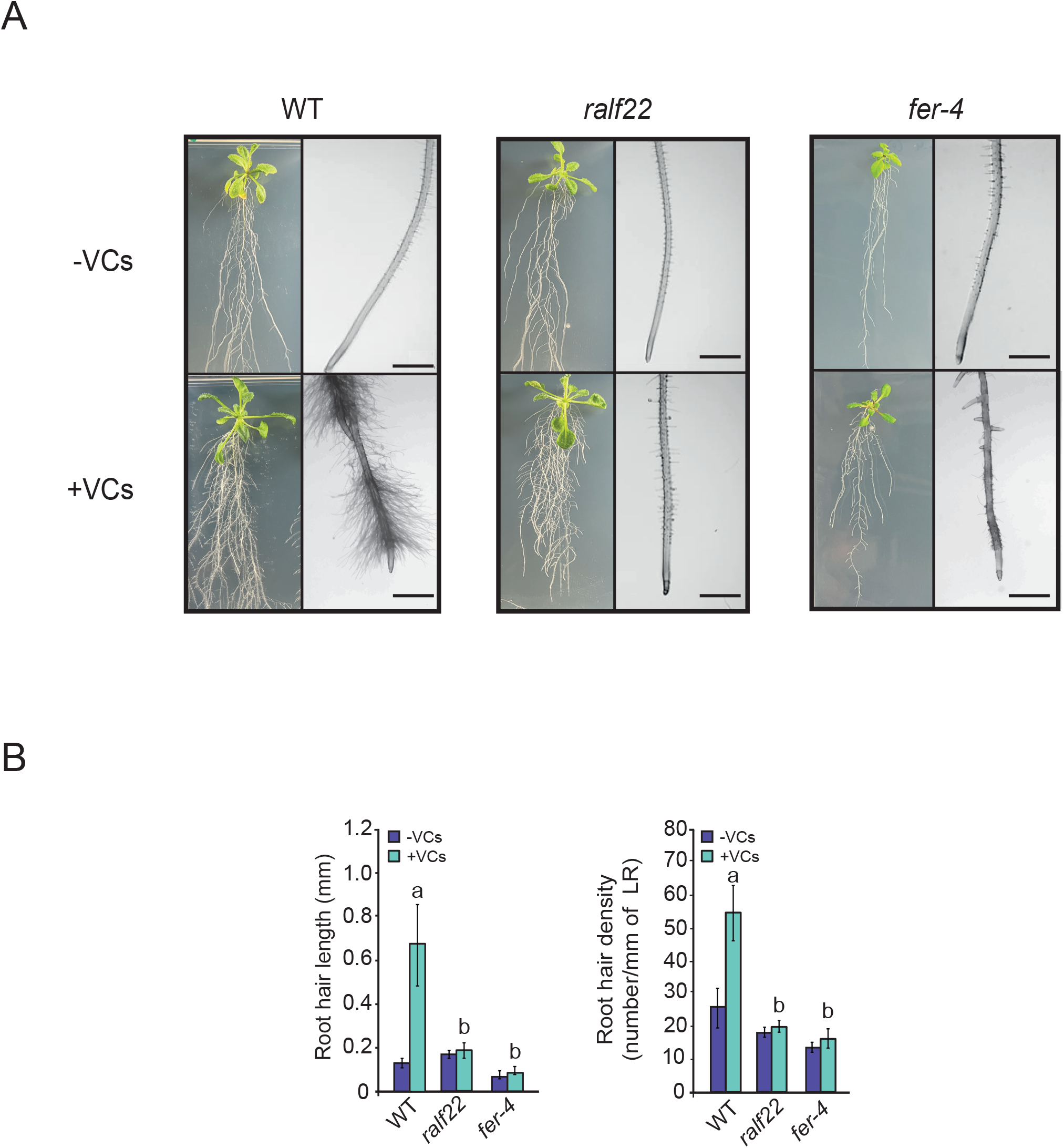
(A) External phenotypes and (B) RH parameters (density and length) of WT, *ralf22* and *fer-4* plants cultured on vertical plates in the absence or continuous presence of fungal VCs for one week, with 45 mM sucrose supplementation in the culture medium. Values in panel (B) are means ± SE for three biological replicates (each a pool of 12 plants) obtained from four independent experiments. Letters “a” and “b” indicate significant differences, according to Student‟s t-test (P<0.05), between: “a” VC-treated and non-treated plants and “b” VC-treated WT and mutants. Scale bars in (A), 1 mm.

**Supplemental Table 1.**
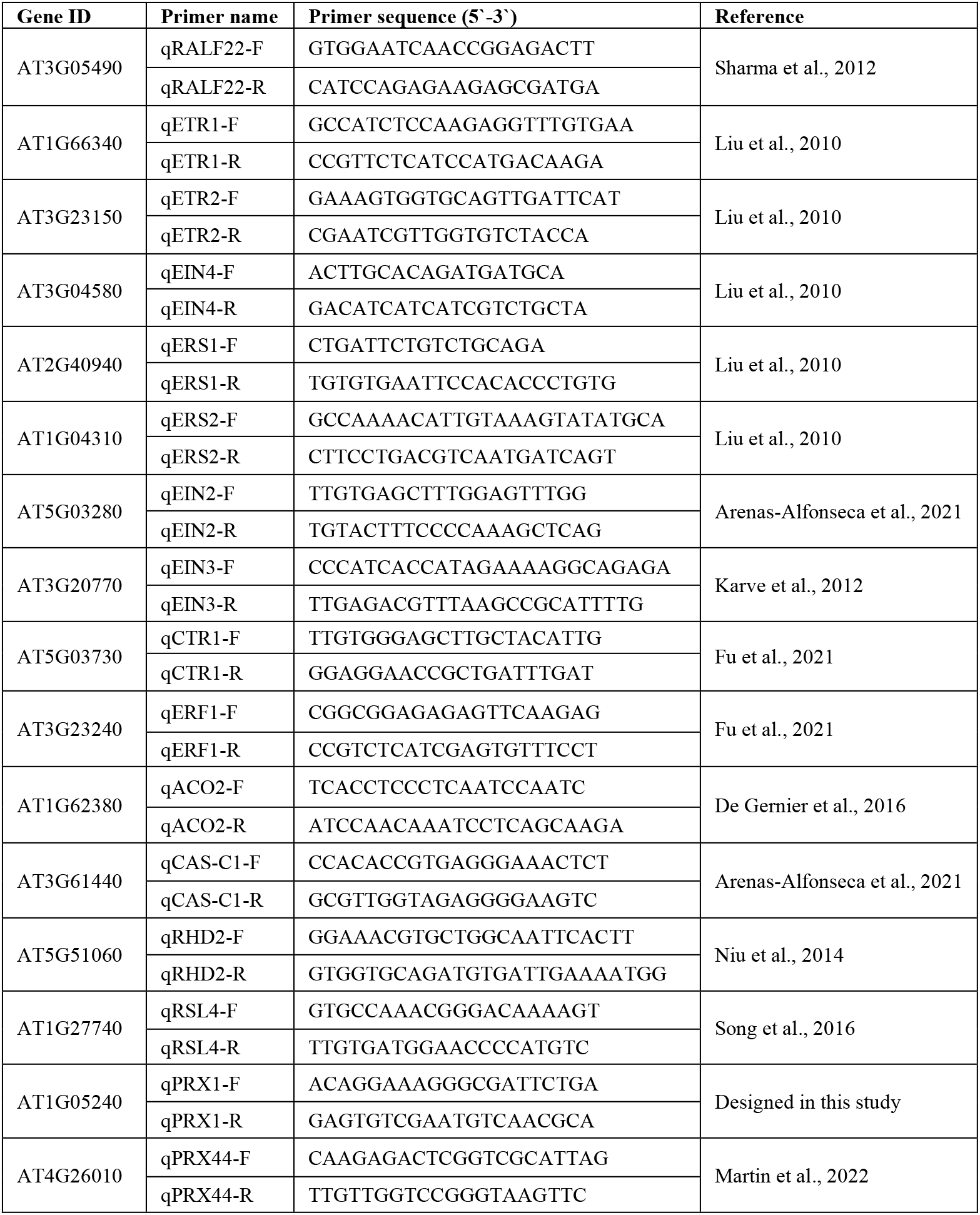
Primers for qRT-PCR used in this study.

